# Eruptive insect outbreaks from endemic populations under climate change

**DOI:** 10.1101/2024.06.27.601089

**Authors:** Micah Brush, Mark A. Lewis

**Affiliations:** Department of Biology, University of Victoria, Victoria, BC, Canada; Department of Mathematics and Statistics, University of Victoria, Victoria, BC, Canada

**Keywords:** Mountain pine beetle, forest pests, outbreak dynamics, endemic populations, climate change

## Abstract

Insects, especially forest pests, are frequently characterized by eruptive dynamics. These types of species can stay at low, endemic population densities for extended periods of time before erupting in large-scale outbreaks. We here present a mechanistic model of these dynamics for mountain pine beetle. This extends a recent model that describes key aspects of mountain pine beetle biology coupled with a forest growth model by additionally including a fraction of low-vigor trees. These low-vigor trees, which may represent hosts with weakened defenses from drought, disease, other bark beetles, or other stressors, give rise to an endemic equilibrium in biologically plausible parameter ranges. The mechanistic nature of the model allows us to study how each model parameter affects the existence and size of the endemic equilibrium. We then show that under certain parameter shifts that are more likely under climate change, the endemic equilibrium can disappear entirely, leading to an outbreak.

## 1 Introduction

Insect outbreaks are often characterized by dramatic year to year changes in population abundances (Barbosa and Schultz 1987). Many species are characterized by eruptive dynamics, where they maintain low densities over a long time before outbreaking at the landscape scale. This pattern includes species such as spruce budworm (Ludwig et al. 1978; Pureswaran et al. 2016), spongy moth (Elkinton and Liebhold 1990), and bark beetles such as spruce beetle (Werner et al. 2006) and mountain pine beetle (Safranyik and Wilson 2006). Outbreaks are therefore particularly hard to predict due to their irregular intervals and dependence on stand conditions and low population densities that are challenging to measure (Berryman and Stark 1985). The impact of insect outbreaks are expected to worsen with climate change (Bale et al. 2002; Logan et al. 2003; Liebhold 2012), creating additional challenges for managers (Day and Pérez 2013; Ayres and Lombardero 2018) and leading to large economic losses (Ayres and Lombardero 2000).

We here focus on the mountain pine beetle (MPB, *Dendroctonus ponderosae* Hopkins) as an example species of these types of endemic to outbreak dynamics. MPB is one of the most destructive forest pests in North America, with outbreaks in the past few decades affecting more than 18 million hectares of forest in Western Canada (Dhar et al. 2016; Corbett et al. 2016). In the historical range in British Columbia, outbreaks were found to occur every 30 to 40 years (Alfaro et al. 2010; Axelson et al. 2009), with evidence that these outbreaks are driven by climatic conditions in combination with forest management (Sambaraju et al. 2019; Safranyik and Wilson 2006).

For MPB, outbreaks often begin from a low level endemic population (Safranyik and Wilson 2006). Beetles in the endemic state preferentially attack trees with compromised defenses from disease, age, or other bark beetles (Safranyik and Wilson 2006; Boone et al. 2011; Bleiker et al. 2014). When conditions are right, beetle populations grow large enough that they are able to mass attack and overcome the defenses of larger, more vigorous trees (Safranyik and Wilson 2006; Raffa and Berryman 1983), increasing their brood size in the higher quality host and thus leading to a transition into an outbreak state.

There are several phenomenological models of this endemic to epidemic transition (Nelson et al. 2008; Cooke and Carroll 2017). These models rely on the idea that weakened and healthy trees each provide an equilibrium state for beetle dynamics. Weakened trees give rise to an endemic equilibrium state at low beetles densities, while healthy trees give rise to an epidemic equilibrium state at high beetle densities, but only when the beetles are able to overcome the defenses of the healthy trees. The existence of both equilibrium states creates bistable dynamics, where the beetle population can transition from the endemic state to the epidemic state if the weakened trees produce enough beetles to overcome the defenses of the healthy trees. This transition could occur if, for example, the beetles are able to increase their brood size in the weakened trees, or if the host defenses of the healthy trees are compromised. Cooke and Carroll (2017) consider their conceptual model in the context of climate change and find that several different impacts of climate change could lead to the disappearance of the endemic equilibrium, allowing MPB populations to suddenly grow to their epidemic state.

Despite the understanding of the biological mechanisms relating to the endemic beetles, there have not been mechanistic models explicitly based on this understanding. Such a mechanistic model could demonstrate how outbreaks occur from endemic populations that are maintained by the existence of low-vigor trees. Given the difficulties in finding and studying low density endemic populations directly in the field (Bleiker et al. 2014), a mechanistic model may be useful in understanding how these endemic populations could change under different environmental conditions.

We here develop a mechanistic model of endemic beetle population dynamics built from first principles. We extend a recent model of MPB (Brush and Lewis 2023) to include low-vigor host trees. This model includes key biological features such as beetle aggregation and a threshold for beetles to overcome host defenses, as well as a forest growth model. We show that this model extension allows for a stable, endemic beetle population, as expected biologically and from the phenomenological models. We then consider realistic ranges for each of the model parameters to investigate under what conditions an endemic beetle population can exist, and how that outcome may be altered under climate change.

## 2 Methods

### 2.1 Model description

We begin with the model described in Brush and Lewis (2023). This model includes key aspects of pine beetle biology (Goodsman et al. 2016) and couples the beetle population dynamics to a forest growth model (Duncan et al. 2015). Beetles in this model aggregate to overcome a host defense threshold, an abstraction of the complex chemical signalling beetles use for mass attack (Raffa and Berryman 1983; Berryman et al. 1985; Safranyik and Wilson 2006). The forest grows with age classes tracking juvenile trees before trees become susceptible when large enough to support beetle populations. New saplings grow wherever there is space on the forest floor.

Mathematically, we use discrete time to represent the annual life cycle of the beetles, with each variable indexed by the year *t*. The variables are defined at the level of a single stand and are the total number of beetles *B*_*t*_ in the stand, the number of susceptible trees *S*_*t*_, the number of infested trees *I*_*t*_, and the number of juvenile trees in each age class *i, j*_*i,t*_. The model parameters are the juvenile survival rate *s*, the number of age classes of juvenile trees before they become susceptible *N*, the host defense threshold *φ*, the beetle brood size *c*, and the beetle aggregation parameter *k*. The beetles are distributed among the susceptible trees according to a negative binomial distribution with aggregation parameter *k* and mean *m*, where the mean number of beetles per tree in year *t* is given by *m*_*t*_ = *B*_*t*_*/S*_*t*_. The probability *p* that *q* beetles are present on a susceptible tree is then given by the probability mass function

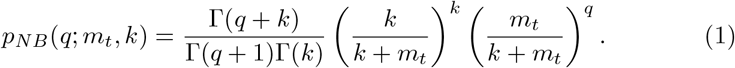

The fraction of surviving trees is given by the function

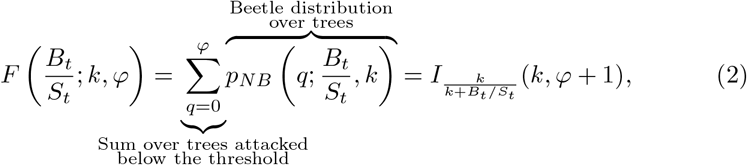

where 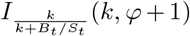 is the cumulative distribution of the negative binomial distribution, which is the regularized incomplete Beta function. For notational simplicity, we drop the explicit dependence on the parameters and write

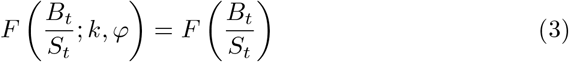

In terms of forest growth, we assume that new growth is light limited, and that infested trees block light for two years, first as red tops and then as grey snags (Safranyik and Carroll 2006; Duncan et al. 2015; Brush and Lewis 2023). We also assume that new saplings replace juvenile trees that do not survive to the next age class. This means the total number of trees *T* is conserved as

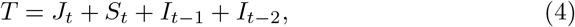

where *J*_*t*_ is the total number of juvenile trees, 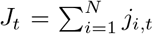.The model can then be written as

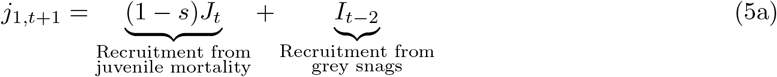

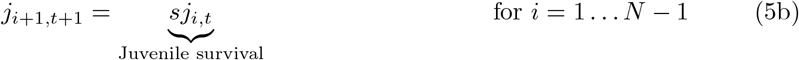

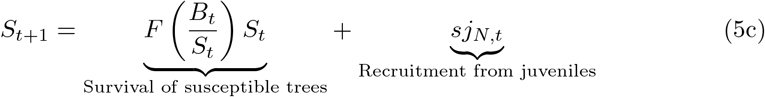

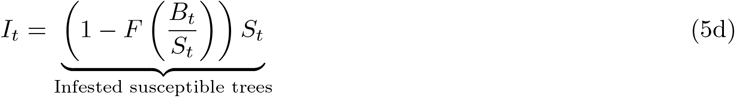

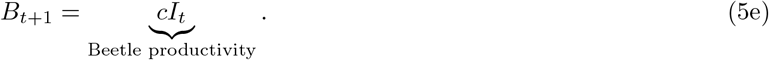

We now extend the model to include low-vigor trees, which represent trees that could be weakened by disease or other pests, older trees with compromised defenses, or trees under more severe drought stress, among other things. These trees will have less resistance to pine beetle attacks, but due to thinner or less nutrient dense phloem will produce fewer beetles once infested (Safranyik and Wilson 2006).

We assume that a small fraction *f* of susceptible trees each year are lowvigor trees. We refer to the remaining fraction 1 − *f* of trees as high-vigor trees. We use the subscript *L* to denote parameters relating to the lower vigor trees and the subscript *H* to denote parameters relating to the high-vigor trees. We denote the threshold to overcome the defenses of these trees as *φ*_*L*_, which should be less than the threshold *φ*_*H*_. Similarly, we denote the brood size in these trees as *c*_*L*_, where again *c*_*L*_ should be lower than *c*_*H*_. Additionally, we assume that the beetles aggregate differently on both high- and low-vigor trees, with parameters *k*_*L*_ and *k*_*H*_. Finally, we assume that a proportion *β* of the available beetles attack low-vigor trees. The mean number of beetles attacking the low-vigor trees is then given by *βB/f S*, or *βm/f* using *m* = *B/S* as before. The mean number of beetles attacking the high-vigor trees is then given by (1 − *β*)*m/*(1 − *f*). Note then that when *β* = *f* and the beetles attack the trees in the same fraction they are present, the mean number of beetles attacking each type of tree simplifies to *m*.

In the analysis, we will allow the fraction *β* to itself be a function of the mean number of beetles present, given evidence that beetles preferentially attack lower vigor trees at low densities and switch to preferentially attack large, higher vigor trees at higher densities (Boone et al. 2011; Bleiker et al. 2014; Safranyik and Wilson 2006). For notational simplicity, we omit this dependence in the following equations.

We turn to the full set of equations including the low-vigor trees. The structure of the juvenile trees is unchanged. We use the same function *F* to describe how beetles overcome tree defenses, but we now have functions with two different mean numbers of beetles, aggregation parameters, and thresholds. We define the function representing the survival of high-vigor trees as

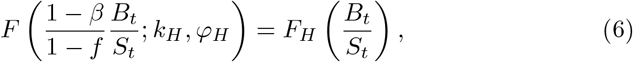

and the corresponding function representing the survival of the low-vigor trees as

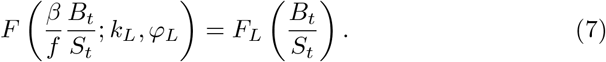

The total number of infested trees is then given by

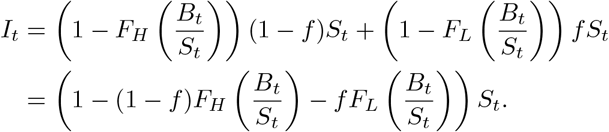

The number of surviving susceptible trees is modified similarly. Note that if *φ*_*H*_ = *φ*_*L*_, *k*_*H*_ = *k*_*L*_, and *β* = *f*, this simplifies to the case without low-vigor trees. The number of emerging beetles is further modified, as the infested lowvigor trees produce a different number of beetles, and is given by

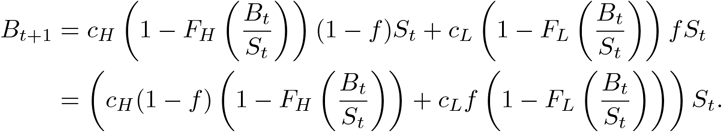

Note that for this equation to simplify to the case without low-vigor trees (i.e. *B*_*t*+1_ = *cI*_*t*_), we additionally require *c*_*H*_ = *c*_*L*_.

The full set of equations is then given by

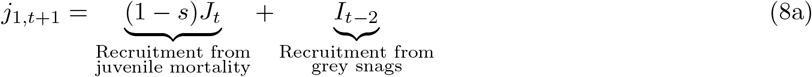

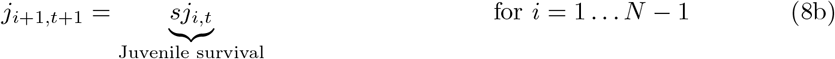

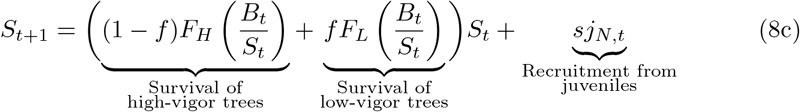

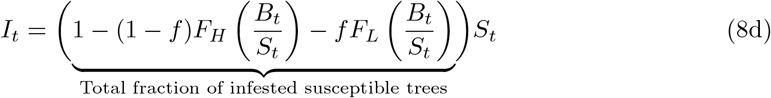

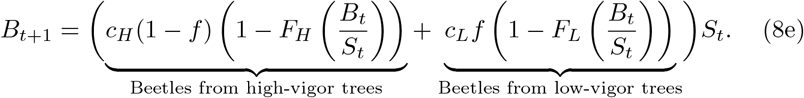

### 2.2 Parameter values

For the high-vigor hosts, parameter values are determined from existing literature as in Brush and Lewis (2025) and Brush and Lewis (2024). Parameter values for the low-vigor hosts are more difficult to determine due to the challenge of collecting data on endemic beetle populations. We therefore allow these parameters to have a wider range of values. The parameter values we consider are summarized in Table 1.

**Table 1:**
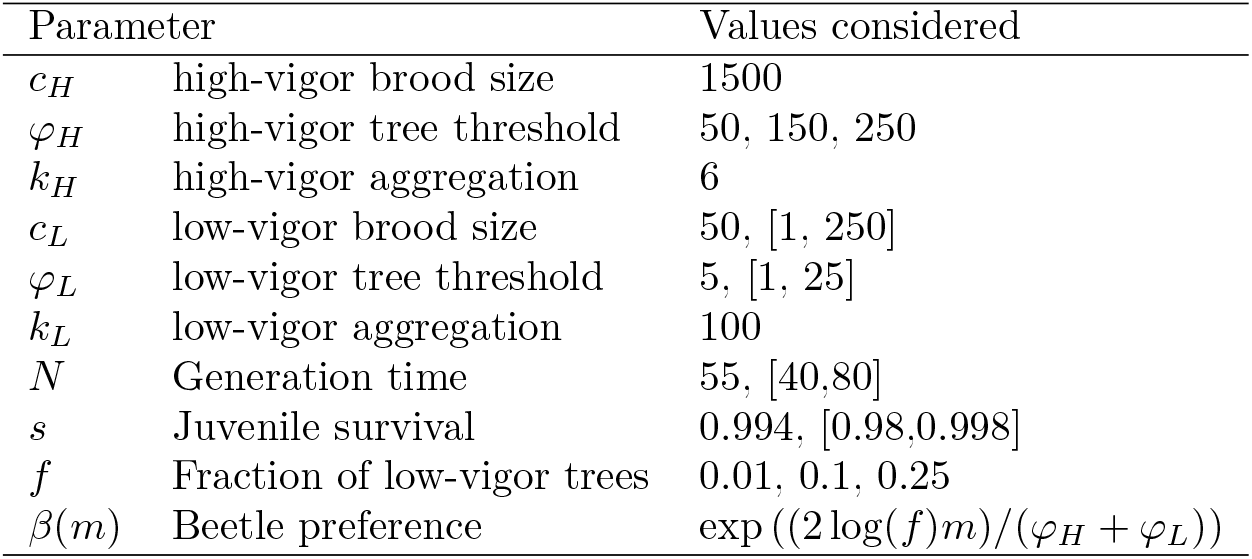
Parameter values for high- and low-vigor trees estimated from the literature. The brood size and tree threshold are in units of beetles per tree. Parameters with a single number are fixed, those with multiple values are varied in discrete increments, and those with a single value followed by values in parentheses are fixed at the single value in some plots and elsewhere varied continuously within the range.

#### Brood size *c*_*H*_

This parameter is likely highly variable given the environment in a given year, as well as non-climate factors such as the size of the host tree (Safranyik and Wilson 2006). We estimate this parameter from historical data (Klein et al. 1978; Raffa and Berryman 1983; Safranyik 1968; Cole and Amman 1969; Safranyik and Wilson 2006), and consider that climate change is likely to increase the average brood size by decreasing overwinter mortality (Régnière and Bentz 2007; Cooke and Carroll 2017). This decrease in overwinter mortality is likely to be realized from fewer very cold winters, as defined by the number of days below -30°C (Hayhoe and Stoner 2019; Eum et al. 2023). In the model extension considered here, we find that the results are not that sensitive to the value of *c*_*H*_, and so we fix *c*_*H*_ = 1500. This value is within the expected historical range and that projected under climate change (Brush and Lewis 2025; Brush and Lewis 2024).

#### Tree threshold, *φ*_*H*_

This parameter is very important for determining the Allee threshold for the beetle population to reach outbreak levels, and so we consider a range meant to capture extreme values that correspond to tree defenses being weakened by increased drought stress under climate change (Safranyik and Wilson 2006; Cooke and Carroll 2017; Eum et al. 2023). Previous work finds a reasonable central value for this threshold in a beetle outbreak to be *φ*_*H*_ = 250 beetles per tree (Raffa and Berryman 1983; Waring and Pitman 1985; Lewis et al. 2010; Peterman 1974; Brush and Lewis 2025). A reasonable lower estimate under current climate conditions is around *φ*_*H*_ = 150 (Brush and Lewis 2025). We additionally allow for the very low value of *φ*_*H*_ = 50, which could reflect a significant decrease in host defenses under frequent or more severe drought.

#### Low-vigor parameters, *φ*_*L*_ and *c*_*L*_

These parameters are very challenging to estimate from data, given the challenge of collecting data on endemic beetles, so we consider a range for both. The threshold *φ*_*L*_ may be only a few beetles per tree for low-vigor hosts (Safranyik and Wilson 2006), as these low threshold trees may be predominately trees already infested by other bark beetles or otherwise have very weakened defenses. We consider a relatively generous range of *φ*_*L*_ = 1 – 25 beetles per tree, though the lower range of this estimate is likely more consistent with available data. We expect that successfully reproducing beetles in low-vigor hosts are likely only able to achieve 1.5 – 2 new beetles per female (Safranyik and Wilson 2006; Klein et al. 1978), with a reasonable maximum of 10 new beetles per female (Raffa and Berryman 1983). Given this, *c*_*L*_ should be at maximum a few times *φ*_*L*_. Again, we consider a generous range from *c*_*L*_ = 1 – 250 beetles per tree, with the most likely values towards the lower end of this range.

#### Beetle aggregation, *k*_*H*_ and *k*_*L*_

We set *k*_*H*_ according to fits to data from Peterman (1974) which indicate that somewhere between 6 and 10 is reasonable (see Brush and Lewis 2025). We note that our results are not particular sensitive to any value in this range and set *k*_*H*_ = 6 here. We set *k*_*L*_ to be very large, *k*_*L*_ = 100, effectively assuming that the beetles are randomly distributed among the low-vigor trees. Biologically, we do this because at very low densities, we do not expect the beetle aggregation pheromones to be very effective, and brood sizes are small, so the beetles should not be very clustered. Note that we are thus effectively replacing the negative binomial distribution with a Poisson distribution for the low-vigor hosts.

#### Tree growth parameters, *N* and *s*

The number of years to become susceptible *N* and the juvenile survival *s* affect the value of the endemic equilibrium and the time dynamics of the outbreak. We explore a reasonable range of these parameters for a well-adapted lodgepole pine stand in Alberta, following Brush and Lewis (2024). We allow *N* to vary between 40 and 80 years, with a central value of 55 years, which comes from site index projections (Monserud et al. 2008) together with a height to dbh model (Huang 1999). We allow *s* to vary between 0.98 and 0.998 1/years, with a central value of 0.994 1/years, based on data from Brown and Navratil (1995), Bedford and Sutton (2000), and Rweyongeza et al. (2007).

#### Fraction of low-vigor trees, *f*

This parameter is likely to vary significantly from stand to stand, as hosts may be compromised from many different causes, such as pathogens, self-thinning, or other bark beetles. Previous work has found that in lodgepole pine, the number of damaged trees suitable for endemic beetles is between 6 – 13 trees per ha (Pokorny 2021). Given a stand density of 1000 to 2000 stems per ha in thinned stands (Baah-Acheamfour et al. 2023), this number of damaged trees corresponds to around 1% of total trees. Other bark beetles have been found in direct association with endemic MPB at similar densities of between 5-10 attacked trees per ha (Safranyik and Wilson 2006). In the case of some pathogens, such as western gall rust, diseased stands may have a very high percent of infested trees (Van Der Kamp and Spence 1987), and, in many stands, the majority of pine are affected by at least one pest (Maclauchlan and Brooks 2022). In stands that are not thinned for forestry, there is also likely to be a supply of low-vigor trees with suppressed growth (Safranyik and Wilson 2006). We here investigate a range between *f* = 0.01 and *f* = 0.25 to represent stands that are likely able to support an endemic beetle population.

#### Beetle preference, *β*

We model *β*(*m*), where *m* is the mean number of beetles per tree *m* = *B/S*, as a decaying exponential. Thus, at low densities the beetles attack only the low-vigor trees, and at high densities nearly all beetles attack the high-vigor trees. This choice of function is made to correspond with observations of beetle behavior (Boone et al. 2011; Bleiker et al. 2014; Safranyik and Wilson 2006). To parameterize the exponential, we assume that at moderate beetle densities the beetles attack the trees in roughly the fraction they are present, so *β*(*m*) = *f* for moderate beetle densities. We choose moderate beetle densities to correspond with the scenario where the mean number of beetles is equal to the average of the two thresholds, *m* = (*φ*_*H*_ + *φ*_*L*_)*/*2. Setting *β*(*m*) = *f* for this value of *m* with the functional form of *β*(*m*) given by a decaying exponential gives

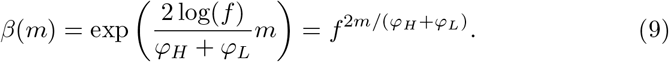

Note that because *f <* 1, log(*f*) is negative, and so (9) is indeed a decaying exponential. The assumption that *β*(*m*) is a decaying exponential parameterized by *β*(*m*) = *f* at an intermediate value of *m* is therefore equivalent to assuming that the beetle preference can be modeled as a power law in *f*.

### 2.3 Numerical simulation under climate change

In addition to our numerical analysis of the fixed points of our model, we conduct a numerical simulation of a beetle outbreak from an endemic population. This simulation is meant to demonstrate how changes in parameters can lead to the disappearance of the endemic equilibrium, and push the system towards the outbreak state. We do this by changing the parameter values from a condition similar to a present day healthy forest to one that could be considered drought stressed, which may be the result of fluctuations in environmental conditions. The numerical values of these parameters are shown in Table 2.

**Table 2:** Parameter values for high- and low-vigor trees varied in our numerical simulation, with their estimated current values and their projected values under climate change. The parameter values are all changed simultaneously by linearly increasing or decreasing from their initial to projected values over 50 years. The brood size and tree threshold are in units of beetles per tree. Parameters not included in this table were held fixed at their values in Table 1.

| Parameter |  | Current value | Projected value |
| --- | --- | --- | --- |
| $c_H$ | high-vigor brood size | 1500 | 1500 |
| $\varphi_H$ | high-vigor tree threshold | 150 | 50 |
| $c_L$ | low-vigor brood size | 50 | 100 |
| $\varphi_L$ | low-vigor tree threshold | 15 | 5 |
| $f$ | Fraction of low-vigor trees | 0.1 | 0.25 |

We begin with the stand at its endemic equilibrium, calculated numerically, and then simultaneously change the parameters in Table 2 from their initial to final values by linearly increasing or decreasing their values from *t* = 10 to *t* = 60. The parameter values therefore change over a 50 year period. Parameters not included in Table 2 are held fixed at their values in Table 1. The parameter changes here are meant to be representative of the kinds of changes we may observe under climate change in the next 50 years, but many combinations of parameter changes would lead to similar results.

## 3 Results

### 3.1 Model fixed points

The mathematical analysis of this model is similar to that in Brush and Lewis (2023). In particular, if *c*_*H*_ = *c*_*L*_ and *β* is fixed rather than a function of *m*, then we can substitute

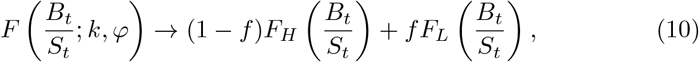

everywhere in the original equations and the analysis follows from Brush and Lewis (2023), but with a more complicated form of the function *F*. If *c*_*H*_ ≠ *c*_*L*_, the model analysis is more complicated as the number of red tops *I*_*t*−1_ cannot be written in terms of the number of beetles in the current year *B*_*t*_, and so we cannot rewrite the right hand side of (8a) in terms of variables at time *t* and must analyze the lagged system.

We now consider the fixed points of the general model defined by (8), where *c*_*H*_ and *c*_*L*_ are independent parameters and *β* is given by (9). The trivial fixed point with no beetles and only susceptible trees (*B* = 0, *S* = *T*) is as in the model defined in Brush and Lewis (2023), and we similarly expect this point to be stable given that the stability of this fixed point was previously shown to not depend on the function *F* (Brush and Lewis 2023). We can solve for the non-trivial fixed points of this model by first dividing both sides of (8e) by *S*, with *m* = *B/S*. The fixed points in *m* are thus given by

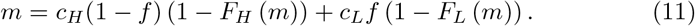

We can then solve for the remaining fixed points by combining (8a) to (8d) together with the conservation of the total number of trees in (4). We set *T* = 1 so we can interpret the number of trees in terms of a fraction of the total stems in the stand. The fixed points, assuming the fraction of infested trees is non-zero, are then given by

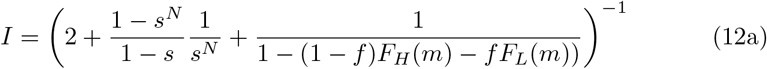

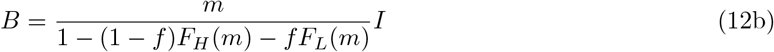

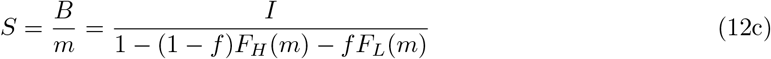

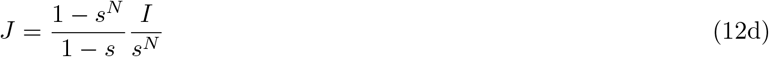

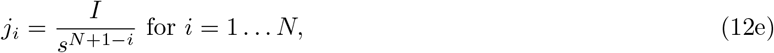

where *m* is calculated numerically from (11).

Under certain parameter conditions, we find four non-trivial fixed points. This is two additional fixed points compared to the previous model. Figure 1 demonstrates graphically how these fixed points arise. The fixed points from (11) are effectively a weighted superposition of those from the model without low-vigor trees, and a model with only low-vigor trees. Without any low-vigor trees present (*f* = 0), we have an Allee effect as before and the upper fixed point is stable. With only low-vigor trees present (*f* = 1), we still have the same model as before with an Allee effect and a stable upper fixed point, but the threshold number of beetles to overcome the Allee effect is much lower, as is the upper fixed point. With only a small fraction of these low-vigor trees then, the associated Allee effect and the fixed point are very small, leading to a small endemic beetle equilibrium, with another higher threshold and fixed point coming from the other susceptible trees.

**Fig. 1.**
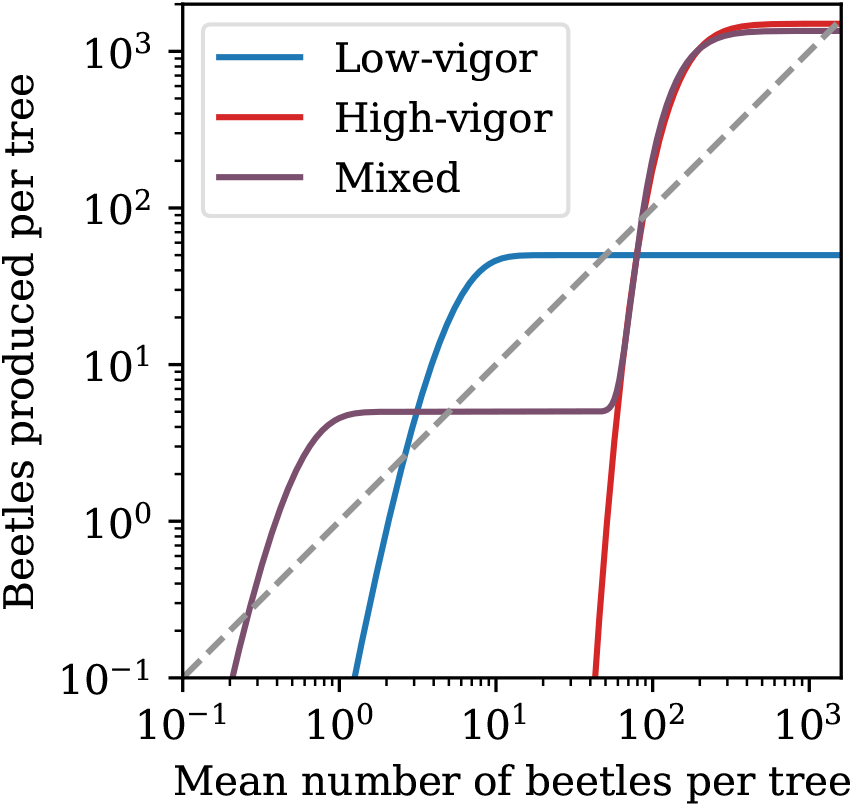
Comparison of the fixed points in the model with no low-vigor trees (*f* = 0), with only low-vigor trees (*f* = 1), and with a small number of low-vigor trees (*f* = 0.1). We fix *φ*_*H*_ = 150, *φ*_*L*_ = 5, and *c*_*L*_ = 50, with other parameters set to their estimated values given in Table 1. With this set of parameters, two additional fixed points appear in the model corresponding to an endemic equilibrium. The fixed points are the points where the solid curves intersect the dashed 1:1 line. The purple line has both high- and low-vigor trees and has four fixed points, and the red line has only high-vigor trees and only two fixed points

Intuitively, given how these fixed points arise, we expect that the trivial fixed point is stable, and the following fixed points alternate between unstable and stable with increasing mean number of beetles (i.e. stable, unstable, stable, unstable, stable, starting from the trivial fixed point). In other words, introducing a small fraction of low-vigor trees is unlikely to change the stability of the fixed points of the model without these trees.

We find that numerical simulations with biologically plausible parameters are consistent with the intuitive stability structure outlined here. In the case *c*_*H*_ = *c*_*L*_ and *β* = *f*, we can calculate the spectral radius numerically by modifying the survival function as in (10). With *c*_*H*_ = *c*_*L*_ = 500, *φ*_*H*_ = 150, *φ*_*L*_ = 5, *f* = 0.1, and other parameters fixed as in Table 1, we find fixed points at *m* = 0, 2.6, 50.2, 97.3, and 494.7 with corresponding spectral radii *ρ* = 0.961, 3.837, 0.989, 2.710, 0.995, which matches our expectations for stability. We note that despite this numerical agreement for these parameter values, there may be small regions of parameter space with unexpected stability behavior, but that these small regions are unlikely to be relevant biologically (as in Brush and Lewis 2023).

Biologically, the additional stable fixed point at low beetle numbers can be interpreted as an endemic beetle population that persists on low-vigor trees, consistent with known behavior (Safranyik and Wilson 2006; Boone et al. 2011; Bleiker et al. 2014). We note that the endemic beetle population comes only from assuming that there is a small fraction of low-vigor trees, that they are easier to infest, and that they have lower brood productivity. We did not assume the existence of endemic beetle populations.

### 3.2 Existence of endemic equilibrium

We now wish to find the conditions and parameter ranges where these additional endemic fixed points exist. We plot the value of the endemic fixed point in Fig. 2 with the parameter values in Table 1. In regions where the endemic equilibrium exists, the value in terms of number of beetles per tree is given by the color. Regions of parameter space where the endemic equilibrium does not exist are shown in black or grey, depending on the reason the endemic equilibrium does not exist. In black regions, no endemic equilibrium exists as the low-vigor hosts are not able to produce enough beetles to achieve a stable equilibrium. In grey regions, no endemic equilibrium exists because the low-vigor hosts produce enough beetles to overcome the higher threshold of the high-vigor trees. Figure 3 demonstrates these different situations graphically, where the black curve is below the 1:1 line until the upper threshold from the high-vigor trees and the grey curve is always above the 1:1 line after passing the lower threshold from the low-vigor trees. The purple curve is included to demonstrate a case where the endemic equilibrium exists, and the values of the parameters are such that we vary only the low-vigor brood size in order to obtain these different behaviors (see the middle left panel in Fig. 2). Biologically, we are particularly interested in the grey regions, as if the parameters of the system shift from a region where the endemic equilibrium exists to a grey region, the beetle population can suddenly overcome the high-vigor threshold and will transition to an outbreak. By contrast, if the parameters of the system shift from a region where the endemic equilibrium exists to a black region, the endemic beetle population is eliminated.

**Fig. 2.**
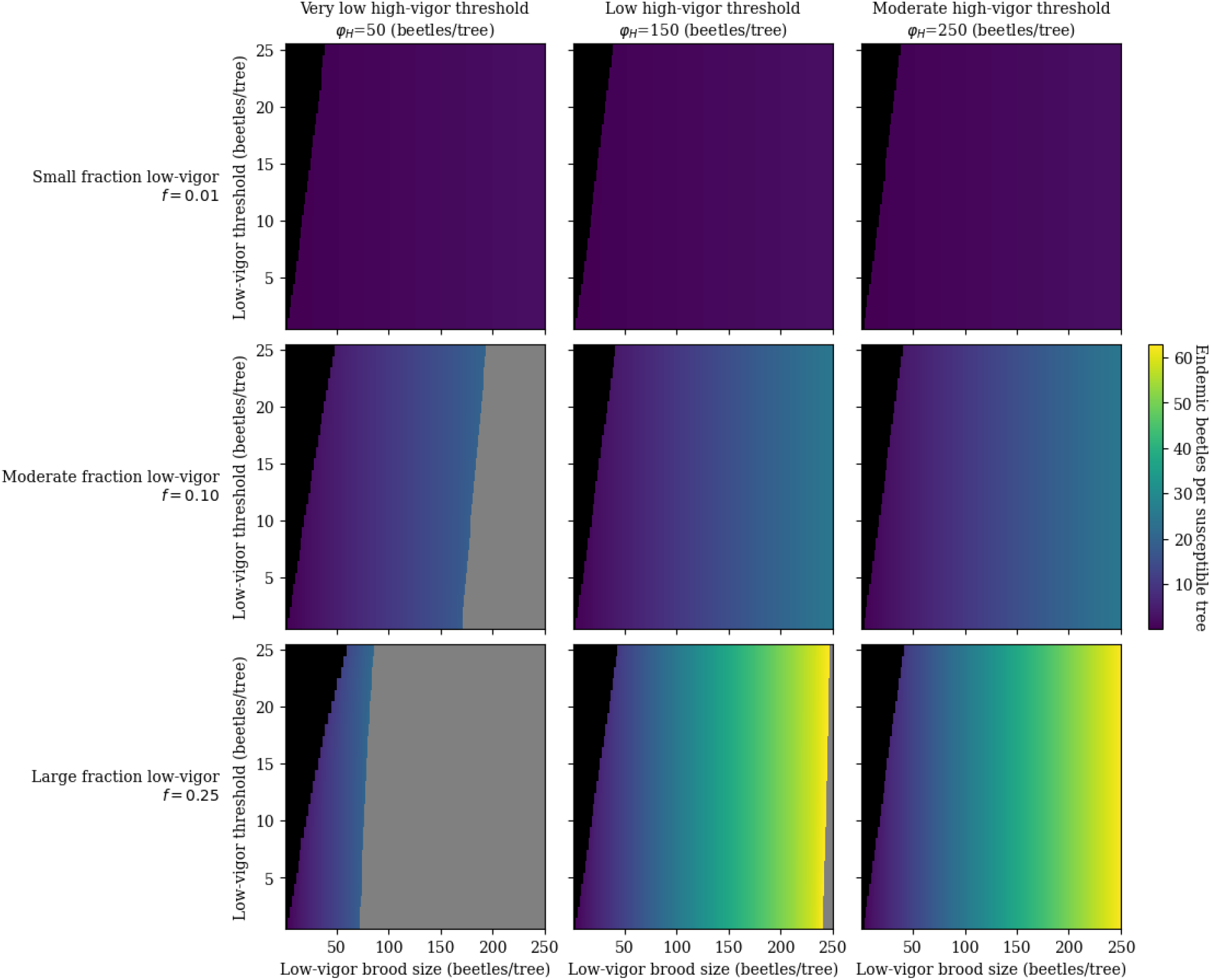
The endemic equilibrium as a function of model parameters. The parameter values are as in Table 1, with rows corresponding to varying the fraction of low-vigor trees *f*, and columns corresponding to varying the threshold number of beetles needed to overcome high-vigor host defenses *φ*_*H*_. Within each plot, we vary the threshold and brood size in the low-vigor trees. The color represents the number of beetles per susceptible tree at the endemic equilibrium. Black regions indicate where no endemic equilibrium exists and the low-vigor hosts do not provide enough beetles to replace themselves (i.e. the beetles produced per tree is below the 1:1 line), and grey regions indicate where no endemic equilibrium exists but the low-vigor hosts produce enough beetles to overcome the Allee threshold for the high-vigor hosts (i.e. the beetles produced per tree is above the 1:1 line, and stays above the 1:1 line with increasing number of beetles). For most realistic values of the model parameters, an endemic equilibrium exists. The grey region, which represents areas where outbreaks could occur from the endemic equilibrium, is largely only present at a very low threshold for the high-vigor hosts

**Fig. 3.**
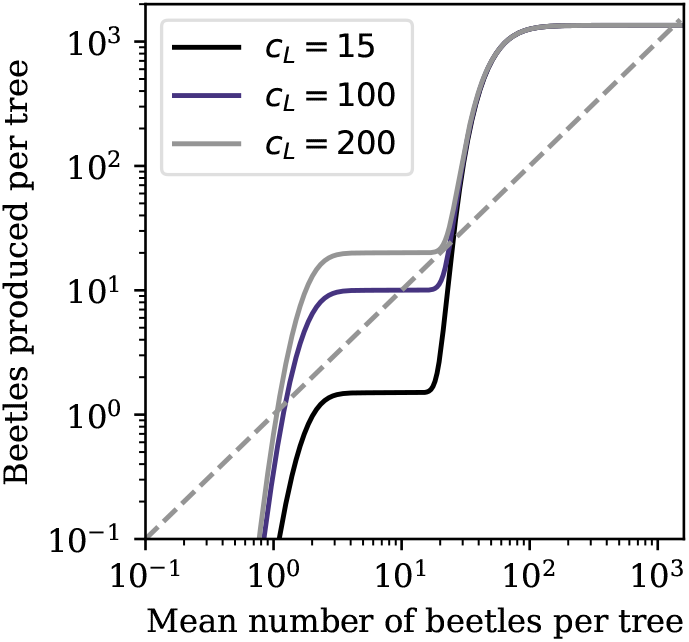
Comparison of the fixed points of the model under different scenarios for the endemic equilibrium. The black curve demonstrates when low-vigor hosts do not produce enough beetles to allow for an endemic equilibrium, and the grey curve demonstrates when the low-vigor hosts produce enough beetles to overcome the threshold of the high-vigor hosts. The purple curve demonstrates when the endemic equilibrium exists. We fix *f* = 0.1, *φ*_*L*_ = 15, *φ*_*H*_ = 50, *c*_*H*_ = 1500, and vary the low-vigor brood size *c*_*L*_ as indicated. This corresponds to the middle left panel of Fig. 2

The regions where the endemic equilibrium does not exist can be roughly approximated analytically in order to gain some intuition for where endemic beetles can persist. As a rule of thumb, an endemic equilibrium is possible if *c*_*L*_ · *f >* Allee threshold of the low-vigor trees. This can be understood as when the beetles are present at low densities, they mostly attack the low-vigor trees (*β*(*m*) ≈ 1, so *F*_*H*_ (*B*_*t*_*/S*_*t*_) ≈ 0), and so, it is possible for the beetles to saturate the survival function corresponding to the low-vigor hosts (*F*_*L*_(*B*_*t*_*/S*_*t*_) ≈ 1) and infest most of these hosts at each time step. Therefore, the number of beetles produced per susceptible tree is approximately *c*_*L*_ · *f* (since *F*_*L*_(*B*_*t*_*/S*_*t*_) ≈ 1 and *F*_*H*_ (*B*_*t*_*/S*_*t*_) ≈ 0). If the value of *c*_*L*_ · *f* is greater than the number of beetles needed to saturate the survival function (i.e. the Allee threshold), then an endemic equilibrium exists. Mathematically, we can see this by first assuming *m* is small such that *β*(*m*) ≈ 1, (9). Then (11) is approximately

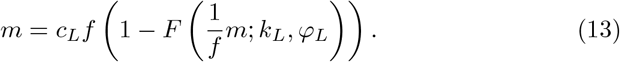

If the right hand side of this expression is greater than the left hand side for some small *m*, then the function passes above the 1:1 line and there will be an endemic equilibrium. Therefore, if there is a (small) value of *m* such that 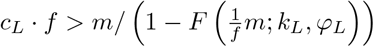, then an endemic equilibrium exists.

We can also look at how each model parameter affects the existence of the endemic equilibrium. The low-vigor parameters determine if the endemic equilibrium exists. The low-vigor tree threshold *φ*_*L*_ controls the rise of the first part of the curve in Fig. 1, and thus whether this curve rises fast enough to be above the 1:1 line. The low-vigor brood size *c*_*L*_ controls the height to which the first part of the curve in Fig. 1 rises, together with the fraction of low-vigor trees *f*. Note that increasing this fraction also decreases the value of the upper equilibrium as there are fewer high-vigor trees with larger brood size. Finally, the high-vigor tree threshold *φ*_*H*_ controls the outbreak Allee threshold.

To connect with data, we need to know the fraction of infested trees at the endemic equilibrium rather than the number of beetles per tree, *m*, as the fraction of infested trees is much easier to measure in the field. In this case, we use (12a) to calculate the fraction of infested trees at the endemic fixed point and plot the results in Fig. 4. (12a) depends explicitly on the tree growth parameters *s*, the juvenile survival, and *N*, the generation time, and so we consider how the fraction of infested trees at equilibrium depends on these parameters. We find that the equilibrium fraction of infested trees does not appear to vary strongly with the beetles parameters of brood size or threshold, and so hold them fixed in this plot.

**Fig. 4.**
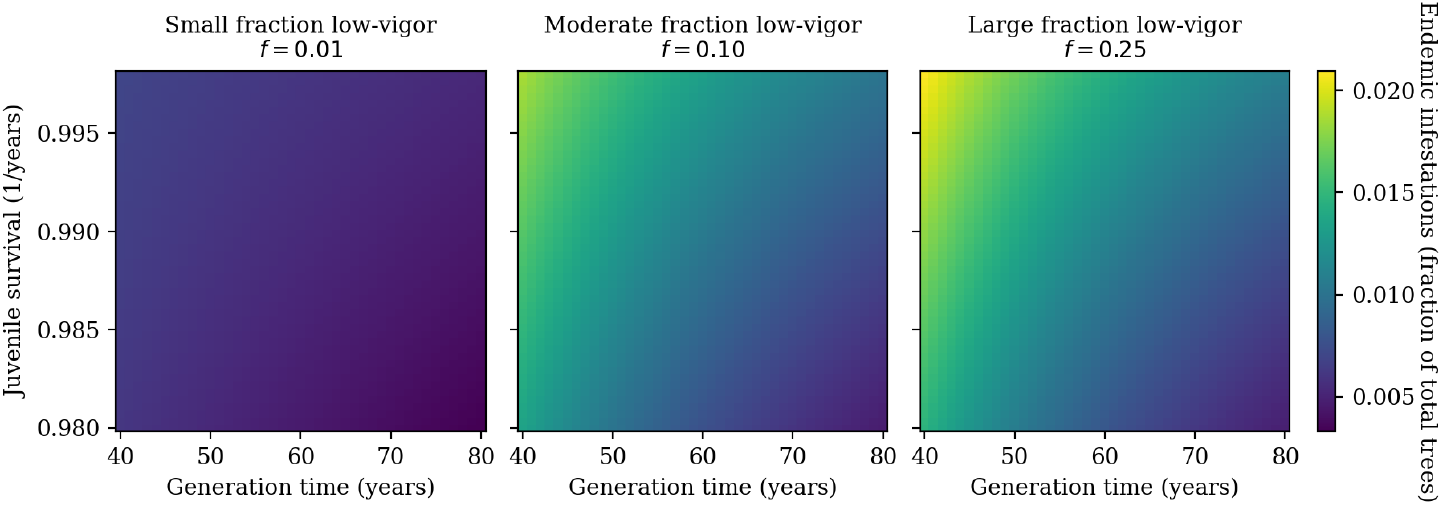
The endemic equilibrium number of infested trees. The color here represents the number of infested trees, in contrast to the equilibrium number of beetles per tree plotted in Fig. 2. Here we vary the parameters *s* and *N*, which enter through the equilibrium number of infested trees, (12a). We consider three different fractions of low-vigor trees *f* and hold other parameters fixed at *φ*_*H*_ = 150, *φ*_*L*_ = 5, and *c*_*L*_ = 50, which is in a parameter space where the endemic equilibrium exists (Fig. 2). The fraction of total trees infested in the endemic equilibrium is in a reasonable biological range, especially in forests with a small fraction of low-vigor hosts

### 3.3 Outbreak from endemic equilibrium

We now turn to our numerical simulation of an outbreak from the lower endemic beetle equilibrium. This outbreak occurs due to the disappearance of the lower endemic equilibrium as the parameter values are altered under climate change. The disappearance of the lower endemic equilibrium can be seen in Fig. 5, with parameters given in Table 2. We can also visualize when this type of outbreak occurs using Fig. 2. If the beetle system is in a region where an endemic equilibrium exists and is at that endemic equilibrium, and the parameter values change such that the system moves to a grey region, the disappearance of the endemic equilibrium will result in an outbreak. Figure 6 shows this visualization for the specific parameters in Table 2, where the orange points represent the location of the system in parameter space.

**Fig. 5.**
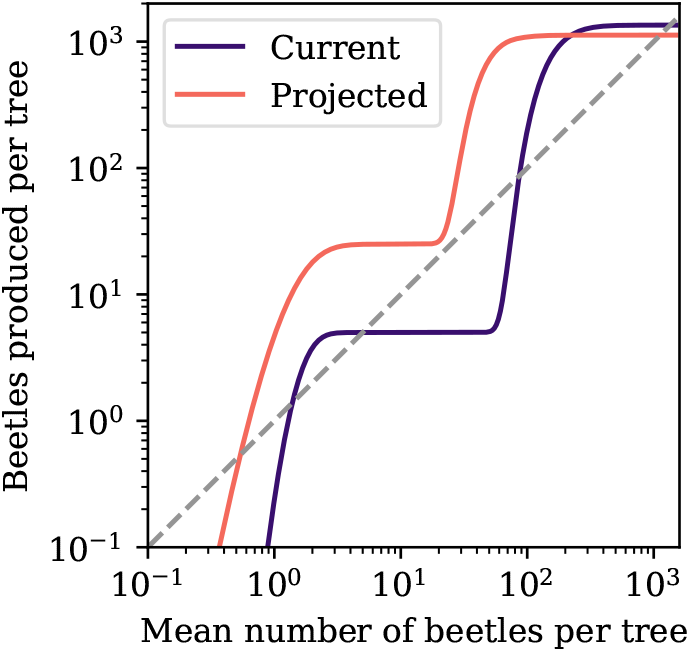
Comparison of the fixed points with parameters set to their estimated current values and their projected values given in Table 2. The endemic equilibrium disappears under the more favourable conditions for beetles, resulting in an outbreak

**Fig. 6.**
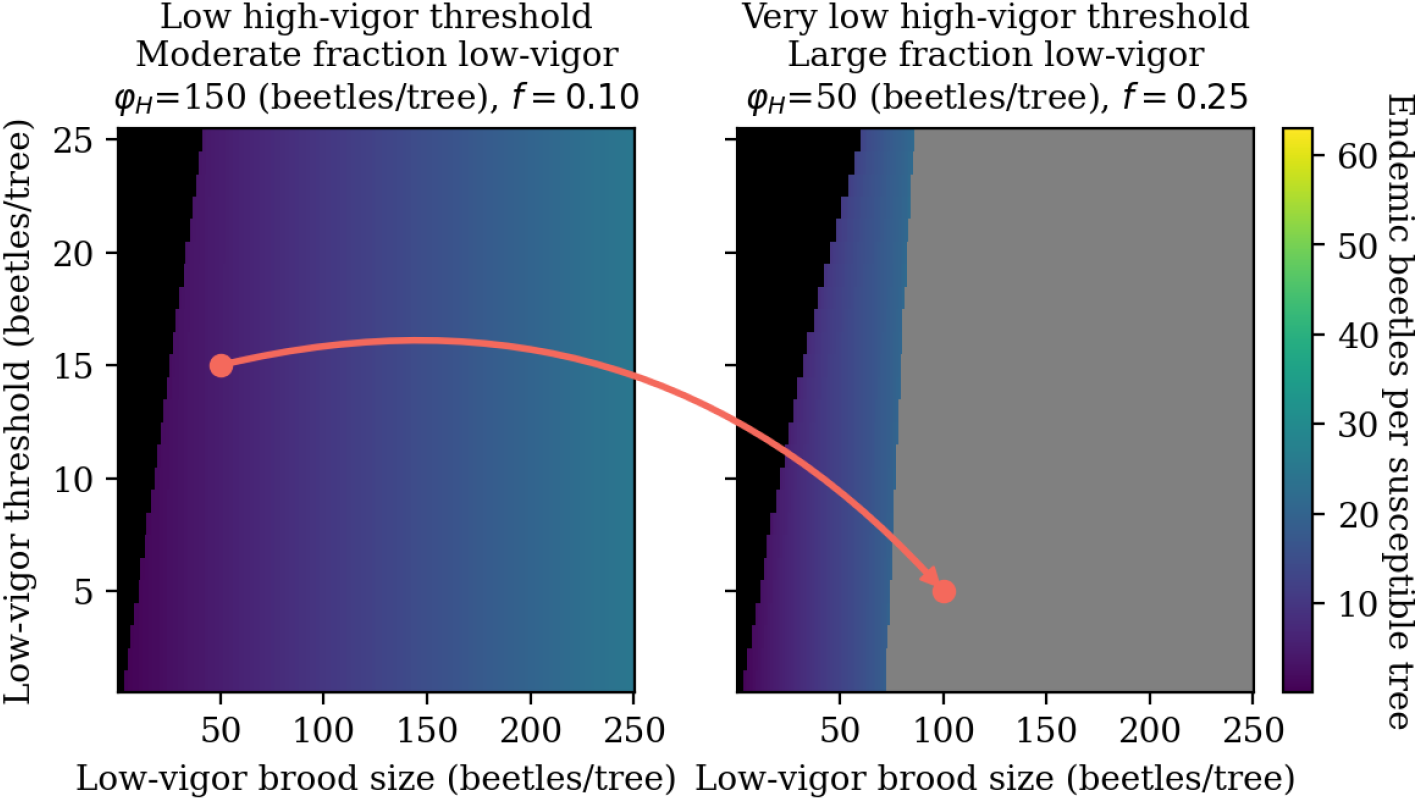
Points show the locations in parameter space for the estimated current and projected values given in Table 2 and shown on plots similar to Fig. 2, with black and grey regions representing areas of parameter space without an endemic equilibrium as in that figure. The orange dots show the location in parameter space before and after the change in parameters, connected with an arrow. This demonstrates how a change in parameters can lead to the disappearance of the endemic equilibrium in the grey region, leading to an outbreak

We plot the resulting outbreak in Fig. 7. As with other simulations with similar versions of this model (Brush and Lewis 2023; Brush and Lewis 2025), the outbreaks are transient with large fluctuations in the beetle populations and with period given by the time of recovery of the forest (*N* + 3). Here, the outbreak begins as a result of the loss of the endemic equilibrium, rather than from the migration of beetles from adjacent stands.

**Fig. 7.**
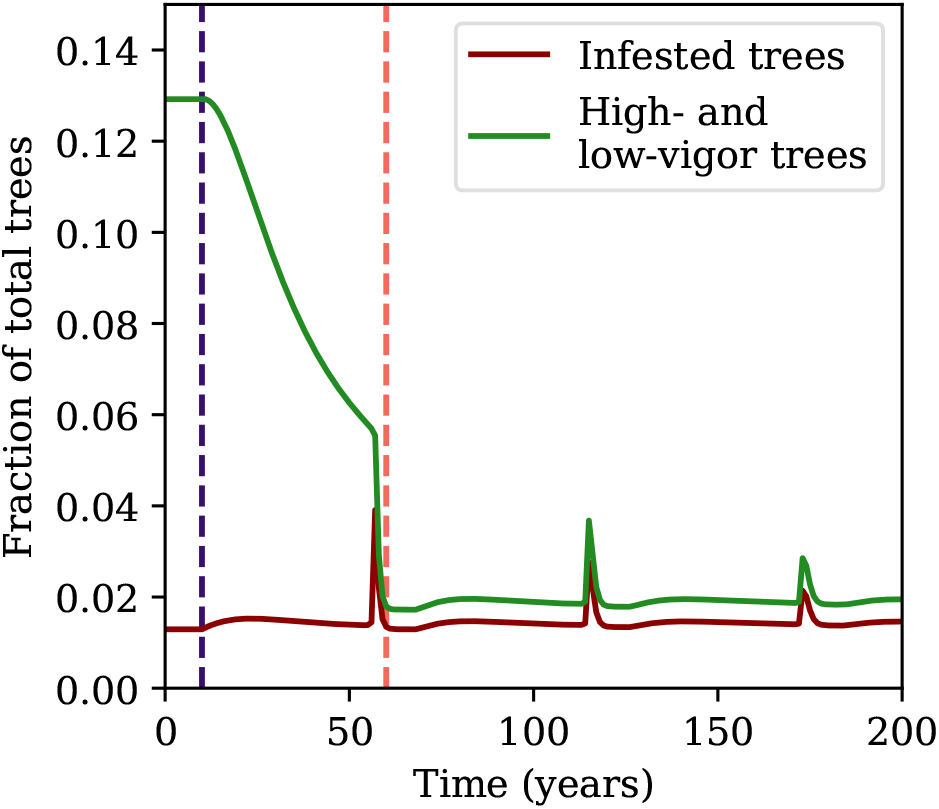
Outbreak from endemic beetles. Infested trees are shown in dark red, and the highand low-vigor trees are combined in green, with both plotted as a fraction of the total number of trees. Note that many trees are juvenile, which are not plotted here. The initial condition is equal to the endemic equilibrium with parameters set to their estimated current values, and the parameters are then reduced linearly over 50 years to their projected values. The first, darker vertical line shows where the parameters begin to change, and the second, lighter vertical line shows where the parameters have attained their final values. The outbreaks are transient pulses of infested trees

## 4 Discussion

We have here extended a mechanistic model of mountain pine beetle population dynamics by adding a small fraction of low-vigor trees. This small fraction gives rise to a small, stable endemic equilibrium, as expected biologically (Safranyik and Wilson 2006) and consistent with previous phenomenological models (Nelson et al. 2008; Cooke and Carroll 2017). In our model, the endemic equilibrium is not assumed but instead arises naturally from the fraction of low-vigor trees, allowing the beetles to reproduce at low densities in weakened trees. The endemic population exists for most realistic values of model parameters (Fig. 2). However, for some regions of parameter space with particularly low high-vigor host resistance, the endemic equilibrium disappears as endemic beetles are able to overcome the threshold for the high-vigor hosts themselves, leading to an outbreak. These parameter regimes are potentially more common with climate change, which would make it more likely for outbreaks from endemic to epidemic conditions.

The number of infested trees at the endemic equilibrium predicted from the model is realistic compared to data. Note that while the model parameters were estimated from data, the number of infested trees at the endemic equilibrium is a model prediction, implying that our model is correctly capturing aspects of mountain pine beetle dynamics. From Fig. 4, we find that between 0.5 and 2% of total trees are infested at the endemic equilibrium. Given stand densities of 1000 to 2000 stems per ha for thinned commercial stands (Baah-Acheamfour et al. 2023), this fraction corresponds to 5 to 40 endemic infested trees per ha. Endemic populations are frequently considered to be stands with less than 5 attacked trees per ha (Boone et al. 2011), with studies finding between 3 – 6 infested trees per ha in mature lodgepole pine stands (Safranyik and Wilson 2006) and 1 – 2 attacked trees per ha in the region of Alberta where lodgepole and jack pine hybridize (Bleiker et al. 2014). Another study fitting a model in British Columbia found 7 – 14 attacking beetles per ha (Koch et al. 2021), which for *φ*_*L*_ = 5 roughly corresponds to 1 – 2 infested trees per ha. These estimates from available data correspond to the low end of our model predictions for the endemic equilibrium, and specifically are attained for a small fraction of low-vigor trees of around 1%. This value is also perhaps the most realistic for endemic populations given available data (Pokorny 2021; Safranyik and Wilson 2006), but we note that this fraction may increase during the transition from endemic to outbreak population levels.

This model provides a mechanistic underpinning for the phenomenological models of Nelson et al. (2008) and Cooke and Carroll (2017). These models also assume that low-vigor trees give rise to an additional fixed point. In Nelson et al. (2008), low-vigor hosts give rise to a second peak in the recruitment curve and thus an endemic equilibrium. In Cooke and Carroll (2017), recruitment is modeled as the combination of epidemic and endemic recruitment curves. The epidemic recruitment curve comes from assuming an increasing growth rate at low density as beetles are able to overcome host defenses (Boone et al. 2011) followed by a decreasing growth rate at high density as beetles saturate the available trees (MacQuarrie and Cooke 2011). The endemic recruitment curve assumes beetles are able to take advantage of stressed trees at low densities, but the beetle growth rate decreases with density given competition for a limited number of stressed trees.

Cooke and Carroll (2017) also consider the effects of climate change on the endemic niche and the potential for outbreaks. Using their model, they consider three perturbations which may lead to a saddle-node bifurcation whereby the endemic niche disappears and the endemic beetles transition to outbreak levels. The three scenarios they consider are climate warming leading to increased brood size from increased overwinter survival, which here corresponds to an increase in *c*_*L*_ and *c*_*H*_, host defense weakening through increased drought stress, which here corresponds to a reduction in *φ*_*H*_, and forest degradation, which here corresponds to an increase in *f*. In all cases, we find that these changes in the parameters of our model can also lead to an outbreak from the endemic equilibrium (Fig. 2). In particular, our simulation shown in Fig. 7 corresponds to a scenario where all three of these changes occur, which leads to the exact scenario discussed in Cooke and Carroll (2017).

Compared to the original model without low-vigor trees (Brush and Lewis 2023), this model extension allows for more realistic outbreaks from endemic beetles rather than from immigrating beetles. In the original model, beetles immigrating from an adjacent stand were required to overcome the Allee threshold and drive the system to transient outbreaks, but here outbreaks can occur directly from changing parameter values leading to the disappearance of the endemic equilibrium, as in Fig. 7. Biologically, this is often how mountain pine beetle outbreaks begin (Safranyik and Wilson 2006). Another difference from the original model (Brush and Lewis 2023) is that the dynamics in the model presented here depend directly on the parameter *φ*_*H*_ rather than on the ratio *φ*_*H*_*/c*_*H*_. This difference in parameter dependence arises from the fact that our model has two brood size parameters, *c*_*H*_ and *c*_*L*_, and so we can no longer nondimensionalize by the brood size in the high-vigor trees (i.e. we can no longer divide through by *c*_*H*_ in (8e)). Other aspects of the outbreak dynamics under climate change that are not dependent on the endemic beetle dynamics are studied in Brush and Lewis (2024).

Our model has several limitations. Most importantly, parameter uncertainty is quite high. We have tried to mitigate this uncertainty by considering a wide range of parameter values, and we find that not all parameter values lead to realistic levels of endemic infestation. Our projected parameter values under climate change are particularly uncertain given the complicated impacts of climate change on pine beetle dynamics (Cooke and Carroll 2017; Bentz et al. 2022). A separate limitation of our model, unrelated to parameter uncertainty, is that our model produces an unrealistic behavior at the endemic equilibrium. For the endemic equilibrium to be in balance, beetles consume all of the available low-vigor hosts in each year. The low-vigor hosts are then replaced in the following year. This behaviour is unrealistic as very small numbers of beetles are unlikely to be able to find and successfully infest all possible low-vigor hosts. That beetles are unlikely to infest all low-vigor hosts is consistent with the available data, as estimates of the number of available low-vigor hosts range from 6 – 13 trees per ha (Pokorny 2021), while endemic population sizes are slightly lower at around 5 trees per ha (Boone et al. 2011; Safranyik and Wilson 2006; Bleiker et al. 2014). However, the number of infested trees at the endemic equilibrium that we find here are still within a biologically plausible range. Given that, we can interpret the fraction *f* of low-vigor hosts as the effective fraction of low-vigor hosts that are available to the endemic beetle population.

Future work could include more detailed simulations. For example, we could draw parameter values from distributions rather than hold them fixed to address some degree of parameter uncertainty, or we could add immigrating beetles in addition to the endemic beetles. We could also make this model spatially explicit, as in Brush and Lewis (2025). We additionally note that there are many possible reasonable assumptions for the functional form of *β*(*m*), and the choice we have made here is certainly not unique. For example, we could try a different functional form such as a sigmoidal function to attempt to model when the beetles switch from attacking low-vigor to high-vigor hosts.

In summary, we find that realistic values of our model parameters lead to biologically plausible levels of endemic infestations. We find that a low fraction of low-vigor trees, together with a small low-vigor brood size and low-vigor threshold, yields observed levels of endemic infestations. Given that a wide range of parameter values allow for an endemic equilibrium, our model suggests lodgepole pine forests in Alberta are likely to enter endemic states following the most recent outbreaks. Under climate change, model parameter values may shift towards values that lead to a higher risk of undergoing an endemic to outbreak transition. More generally, given the difficulty of monitoring low-level endemic populations of forest pests, mechanistic models such as this one can be useful for understanding these endemic populations and how they may change in the future.

## Statements and Declarations

### Data availability statement

Data sharing is not applicable to this article as no datasets were generated or analysed during the current study.

## Acknowledgments

We would like to thank Evan Johnson for his feedback on this project. We would also like to thank all members of the Lewis Research Group and TRIA-FoR for their support and feedback. We thank two anonymous reviewers for their feedback. MAL gratefully acknowledges the Gilbert and Betty Kennedy Chair in Mathematical Biology. Funding for this research has been provided through grants to the TRIA-FoR Project to MAL from Genome Canada (Project No. 18202) and the Government of Alberta through Genome Alberta (Grant No. L20TF), with contributions from the University of Alberta and fRI Research (Project No. U22004). This work was supported by Mitacs through the Mitacs Accelerate Program, in partnership with fRI Research. MB acknowledges the support of the Natural Sciences and Engineering Research Council of Canada (NSERC), [PDF – 568176 - 2022].

